# Intergenerational epigenetic signatures of prenatal adversity and their role in emerging child psychopathology

**DOI:** 10.64898/2026.07.20.739628

**Authors:** Anna Constantino-Pettit, Alexandre A. Lussier, Mia Ruppel, Erin C. Dunn, Darina Czamara, Tara Smyser, Ryan Bogdan, Barbara Warner, Christopher Smyser, Deanna Barch, Cynthia Rogers, Joan Luby

## Abstract

**Background:** Adversity during pregnancy is associated with alterations in offspring brain development. DNA methylation (DNAm) is a type of epigenetic modification is a putative mechanism for the intergenerational transfer of prenatal adversity on developmental outcomes. We examined the effects of prenatal social disadvantage (PSD) and prenatal psychosocial stress (PSS) on offspring epigenome-wide DNAm at four time points from infancy to age 4, and tested whether persistent DNAm signals mediated associations between PSD, PSS, and child psychopathology at ages 4-6.

**Methods:** Longitudinal DNAm data (birth, Y1, Y2, Y3) was derived from salivary tissue of 281 infants (43.7% female) in the eLABE study. We examined epigenome-wide associations between DNAm and PSD and PSS. Linear models were adjusted for age, child sex, child race, maternal tobacco smoking, cell type composition, and batch effects. Mediation analyses tested whether birth DNAm mediated associations between prenatal exposures and internalizing and externalizing symptoms at ages 4-6.

**Results:** PSD was associated with 47 FDR-significant CpGs at birth and 3 at year 1. PSS was associated with 3 FDR-significant CpGs at birth. Thirteen CpGs showed PSD-associated significance across all four timepoints, including two CpGs associated with genes involved with neuronal maturation (*BCL11B*) and one CpG associated with a gene implicated in brain vascular health (*ZNF474).* Exploratory mediation analyses revealed an indirect effect of methylation at *ZNF474* (cg19980369; *p*=0.028; *p*_FDR_=0.360) on the association between PSD and year 4-6 externalizing symptoms.

**Conclusions:** PSD was associated with epigenetic signatures at birth, with a subset of associations persisting across early childhood and converging on cellular stress response biology. PSS showed minimal epigenetic associations, suggesting differential biological embedding of structural versus psychological dimensions of adversity.

## Introduction

Social Determinants of Health (SDoH), encompassing access to basic necessary resources, quality education, healthcare, neighborhood safety, and economic stability, have been shown to negatively affect both mothers’ health during pregnancy^1^ and early neurodevelopment in their offspring^2–4^. Adverse SDoH are particularly impactful for both mother and baby from the time of conception through the first years of life. Pregnancy is a sensitive period for fetal neurodevelopment^5^, with tremendous brain growth occurring prenatally through preschool age^6^. As a result, environmental influences during this sensitive period may lead to enduring changes in brain growth and development.

Prenatal adverse experiences, particularly those related so social disadvantage (PSD), have been associated with alterations in offspring brain structure and function evident at birth. Neonates born to mothers facing greater socioeconomic disadvantage during pregnancy show reductions in cortical, subcortical, and cerebellar gray matter volumes^7 8 9 10^, differences in white matter microstructure^11 12^, and altered functional connectivity^13^. These differences in neonatal brain functional connectivity have also been found to mediate the association between prenatal social disadvantage and cognitive and socioemotional outcomes in early childhood^14^. However, it is less clear how different biological mechanisms facilitate this intergenerational association between prenatal adversity and newborn brain development. However, epigenetic changes have been proposed as one possible mechanism. For instance, Nelson & Gabard-Durnam’s conceptual model of Sequelae of Adverse Experiences^15^ identifies DNA methylation (DNAm) as a mechanism of biological embedding from which downstream impaired brain development and dysregulated stress reactivity may follow. Similarly, Monk et al.’s Mechanisms and Pathways of Maternal Distress and Fetal and Child Brain-Behavior Development conceptual framework posits that DNAm of glucocorticoid-related genes (e.g., *FKBP5, NR3C1*) affects regulation of stress-response pathways to adversely influence fetal brain programming and, ultimately, child psychopathology^16^.

### Epigenetics as a Mechanism of Intergenerational Transfer

Epigenetic modifications are a plausible candidate mechanism for these intergenerational associations. Capable of influencing gene activity without altering the underlying DNA sequence itself, epigenetic modifications are a key upstream mechanism that may result in differences in gene *expression*. DNAm, one type of epigenetic modification, involves the addition of chemical markers (methyl groups) directly to the DNA structure. These chemical markers bind primarily to locations in the DNA structure where a cytosine nucleotide (C) is directly followed by a guanine nucleotide (G), linked by a single phosphate bond (p; thus, ‘CpG’ sites). Once methylated, and particularly in regulatory gene regions such as promoter regions, it is more difficult for DNA to be transcribed into RNA. This may ultimately result in gene silencing. DNAm is uniquely positioned to be a mechanism of intergenerational transmission for three reasons. First, it is modifiable, with DNAm in both human^17^ and animal^18^ studies demonstrating responsivity to environmental inputs. Second, it is persistent, with changes in DNAm being shown to be preserved both across the lifecourse and intergenerationally^19^. Third, indices of DNAm changes have been associated with psychopathology in adolescence^20^.

### Prenatal Factors, Epigenetic Signatures, and Early Childhood Psychopathology

The literature linking prenatal adversity to offspring DNAm is still emerging. Recent large-scale efforts have linked prenatal stress with offspring DNAm: Ruehlman et al. recently identified a convergent methylation signal across twelve cohorts in the Pregnancy and Childhood Epigenetics (PACE) consortium at cg26579032 (*ALKBH3*)^21^. However, this study (along with the majority of offspring DNAm studies to date) rely on a single timepoint. Serial ascertainment of early childhood DNAm during neuroplastic periods allows for an examination of the timing of DNAm responsivity in relation to prenatal adversity. It is also largely unknown whether the epigenetic signature of prenatal adversity is stable across the early childhood window from infancy through preschool. Other groups have found evidence that different types of adversity are associated with unique DNAm signatures, whether this be within a single generation^22^ or intergenerationally^23^. Finally, no studies to date have examined the entire intergenerational pathway from exposure (prenatal adversity) to outcome (child psychopathology) with epigenome-wide DNAm in early childhood as a mediator. A recent research review called for more studies to examine this pathway in its entirety^24^.

### The Present Study

The current study begins to address some of these gaps. We leverage the Early Life Adversity, Biological Embedding, and Risk for Developmental Precursors of Mental Disorders (eLABE)^25^ cohort, which is a longitudinal sample of mother-infant dyads with deep characterization of both maternal and child phenotype, and child DNAm with yearly sampling from birth through age three. We conducted an epigenome-wide association study (EWAS) of two dimensions of prenatal adversity, prenatal social disadvantage (PSD) and prenatal psychosocial stress (PSS), on child DNAm at each timepoint. We then conducted a mediation analysis with top loci from the EWAS in which we examined child DNAm as a mediator in the relationship between prenatal adversity and early childhood psychopathology at age four-to-six.

By characterizing the developmental epigenetic landscape of two prenatal exposure types across four timepoints, this work aims to clarify both the mechanism and the timing of biological embedding in early life, informing both our understanding of one way prenatal adversity shapes child neurodevelopment and pointing to possible windows during which intervention may be most consequential.

## Methods and Materials

### Participants

Participants were drawn from the eLABE cohort (n=398 mother-infant dyads). Pregnant mothers were recruited from a Midwestern academic medical center during the first trimester of pregnancy, oversampled for poverty, and followed longitudinally. Exclusion criteria for eLABE included multiple gestation pregnancies, infections associated with congenital disease, alcohol use, and illicit drug use (tobacco and cannabis use was not excluded). The current analysis utilized a subset of eLABE mothers whose children had DNAm data collected at one or more of four cross-sectional timepoints: birth (n = 215), year 1 (n = 189), year 2 (n = 130), or year 3 (n = 169). The year 2 collection timepoint and the beginning of the year 3 collection timepoint coincided with the COVID-19 pandemic and resulted in lower total data collection compared to the other 3 timepoints. All study procedures were approved by the Washington University Human Research Protection Office (IRB #201703145) and written informed consent was obtained from mothers at enrollment for both their own and their child’s participation.

### Prenatal Exposures

The two main exposure variables in the analysis were PSD and PSS, both of which are latent composite variables derived from maternal report during pregnancy that have been widely utilized in prior eLABE analyses^7,13,14^. The PSD composite reflects structural and material aspects of disadvantage (income-to-needs ratio, maternal educational attainment, healthy eating, neighborhood-level area deprivation, and health insurance status). The PSS composite reflects psychological and experiential dimensions of stress (depressive symptoms, perceived stress, experiences of racial discrimination, and lifetime adversity severity and frequency). Composite scores were treated as continuous predictors in all regression models.

### DNAm Quantification, Preprocessing, and Quality Control

Epigenome-wide DNAm from saliva samples collected from infants (Oracollect for Pediatrics, OC-175, DNA Genotek, Inc) and children (Oracollect for Pediatrics, OGR-675, DNA Genotek, Inc.) were collected at each study visit, using the Illumina Infinium Methylation EPICv2.0 BeadChip array. We completed standard data pre-processing and normalization using the *meffil* pipeline^26^.

### Child Psychopathology

Internalizing and Externalizing child psychopathology were derived from the Preschool Age Psychiatric Assessment^27^ administered at the age 4-6 eLABE follow-up visit. We used the internalizing symptom count (range 0–20) and externalizing symptom count (range 0–25).

Symptom counts were treated as continuous given the observed distribution and the use of bootstrapped inference.

### Covariates

Covariates were selected based on best practices within EWAS literature while minimizing collinearity with the exposure composites (e.g., income-to-needs ratio was excluded given its contribution to the PSD composite) and included child race, child sex, maternal tobacco smoking during pregnancy, maternal age at birth, and either gestational age at birth (birth EWAS) or child age (year 1-year 3 EWAS). We estimated the proportion of CD45-positive (immune) cells using *meffil*_1.4.0 and included this as a covariate to account for cell-type heterogeneity in saliva.

Sample plate was also retained as a covariate due to lingering technical batch effects following QC. For the mediation analyses, covariates included child sex, child race, and gestational age at birth.

### Epigenome-wide association studies (EWAS)

We conducted eight EWAS in total. Covariate-adjusted multiple linear regression tested the associations between PSS at four timepoints and PSD at four timepoints. For the EWAS, beta values were converted to *M*-values^28^. Significant CpGs were annotated to genes using the Zhou laboratory EPICv2 manifest. Gene annotation was performed on all significant CpGs using Enrichr^29^. Brain-buccal tissue correlation was evaluated using the IMAGE-CpG database^30^.

Finally, we evaluated methylation quantitative trait loci (mQTLs) to assess genetic confounding.

### Genomic Inflation and Bias Correction

We used a Bayesian approach (Bacon^31^) to adjust for genomic inflation in each EWAS. For each Bacon-corrected model, we applied Benjamini-Hochberg FDR-adjusted p < 0.05.

### Persistence Analysis

We then traced their Bacon-corrected p-values and effect sizes across all four timepoints to examine any persistent DNAm effects over time. Because this analysis was restricted to a pre-specified set of CpGs already identified in primary analyses, we used a relaxed threshold of Bacon-corrected nominal p < 0.05 to flag persistent associations at the year 2 and year 3 timepoints.

### Mediation analyses

We conducted exploratory mediation analysis of top CpGs in the association between PSD, PSS, and year 4-6 internalizing and externalizing symptoms using 1,000 nonparametric bootstrap iterations. For all mediation models, β-values rather than M-values were used as the mediator scale to facilitate interpretability of indirect effects on the DNAm proportion scale. Multiple-testing correction was applied within each outcome family (internalizing vs. externalizing) using the Benjamini-Hochberg FDR procedure across all CpG × exposure tests for that outcome.

### Software and Reproducibility

All analyses were conducted in R version 4.4.1 (2024-06-14) “Race for Your Life”. Primary EWAS regressions used the CpGassoc^32^ package. Mediation analyses used the mediation^33^ package. Reproducible code is stored on the Github repository.

## Results

### Sample Characteristics

Saliva DNAm data were available across four cross-sectional timepoints: birth (n=215), year 1 (n=189), year 2 (n=130), and year 3 (n=169). There were no statistically significant differences in demographic characteristics or exposure distribution across timepoints (PSD Kruskal-Wallis X^2^=3.50, p=0.32; PSS Kruskal-Wallis X^2^=1.51, p=0.68; see **Table 1**). The cohort was majority Black or African American (75 [58%] – 121 [64%] range across timepoints), with an average maternal age at delivery of 29.5 years and mean gestational age at birth of 38 weeks.

**Table 1:**
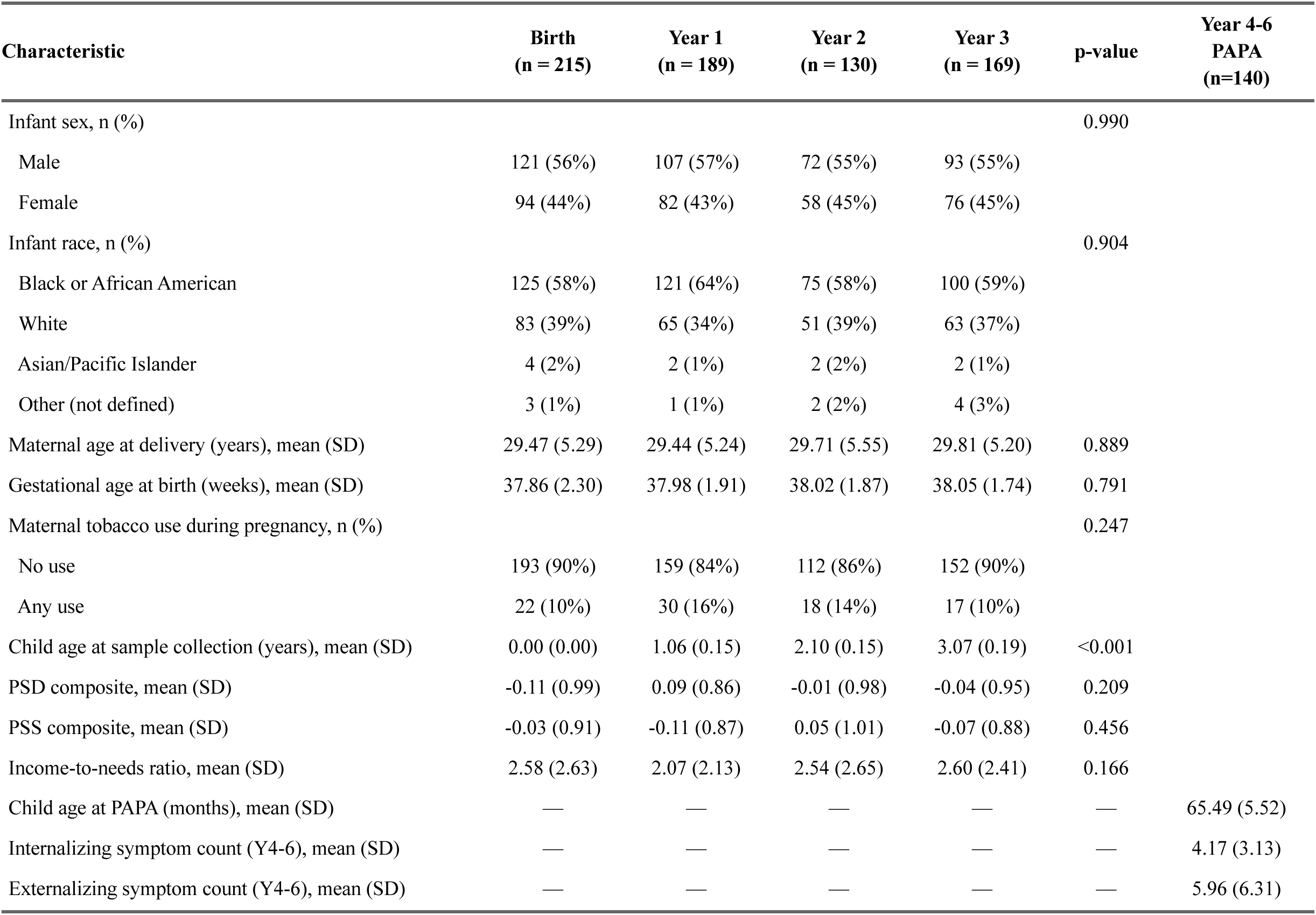
Sample characteristics by methylation collection timepoint. p-values: one-way ANOVA for continuous variables (maternal age, gestational age, child age at collection, PSD, PSS, income-to-needs) and Pearson’s chi-square for categorical variables (infant sex, infant race, maternal tobacco use). Maternal tobacco use is binarized as any use versus none. PSD = prenatal social disadvantage (z-scored); PSS = prenatal psychosocial stress (z-scored). Income-to-needs = household income divided by federal poverty line.

Distributions of the PSD and PSS composites were approximately normal and centered near zero by construction; the two composites were modestly correlated with one another (r=0.24, *p*<0.001).

### Genome-Wide Associations Between Prenatal Exposures and Child Methylation

Eight robust linear regressions tested the associations between the two prenatal exposures (PSD, PSS) and child DNAm at birth, year 1, year 2, and year 3. To account for genomic inflation factors (λ ranged from 1.02 to 1.35), we applied a Bayesian inflation and bias correction approach^31^ to all eight models. Bacon-corrected inflation estimates (σ) ranged from 0.97 to 1.08 for seven of the eight models. The year 3 PSD model produced an elevated bias estimate (μ=+0.45) that we could not reconcile with sample-level characteristics; we therefore hand-calculated the FDR for this model, in line with guidance from the Bacon GitHub documentation.

#### Prenatal Social Disadvantage

PSD was associated with 47 CpGs surviving Benjamini-Hochberg FDR correction (*FDR*<0.05) on bacon-corrected p-values at birth, including three CpGs surviving Bonferroni correction (**Figure 1**). The strongest birth associations were observed at cg09390319 (β=0.26, bacon *p*=6.2 × 10⁻⁹), cg19489172 (β=1.05, bacon *p*=1.1 × 10⁻⁸), and cg01591938 (β=0.34, bacon *p*=9.2 × 10⁻⁸). At year 1, three CpGs survived bacon-corrected FDR, including one that survived Bonferroni correction. There were no significant CpGs at year 2 or 3.

**Figure 1:**
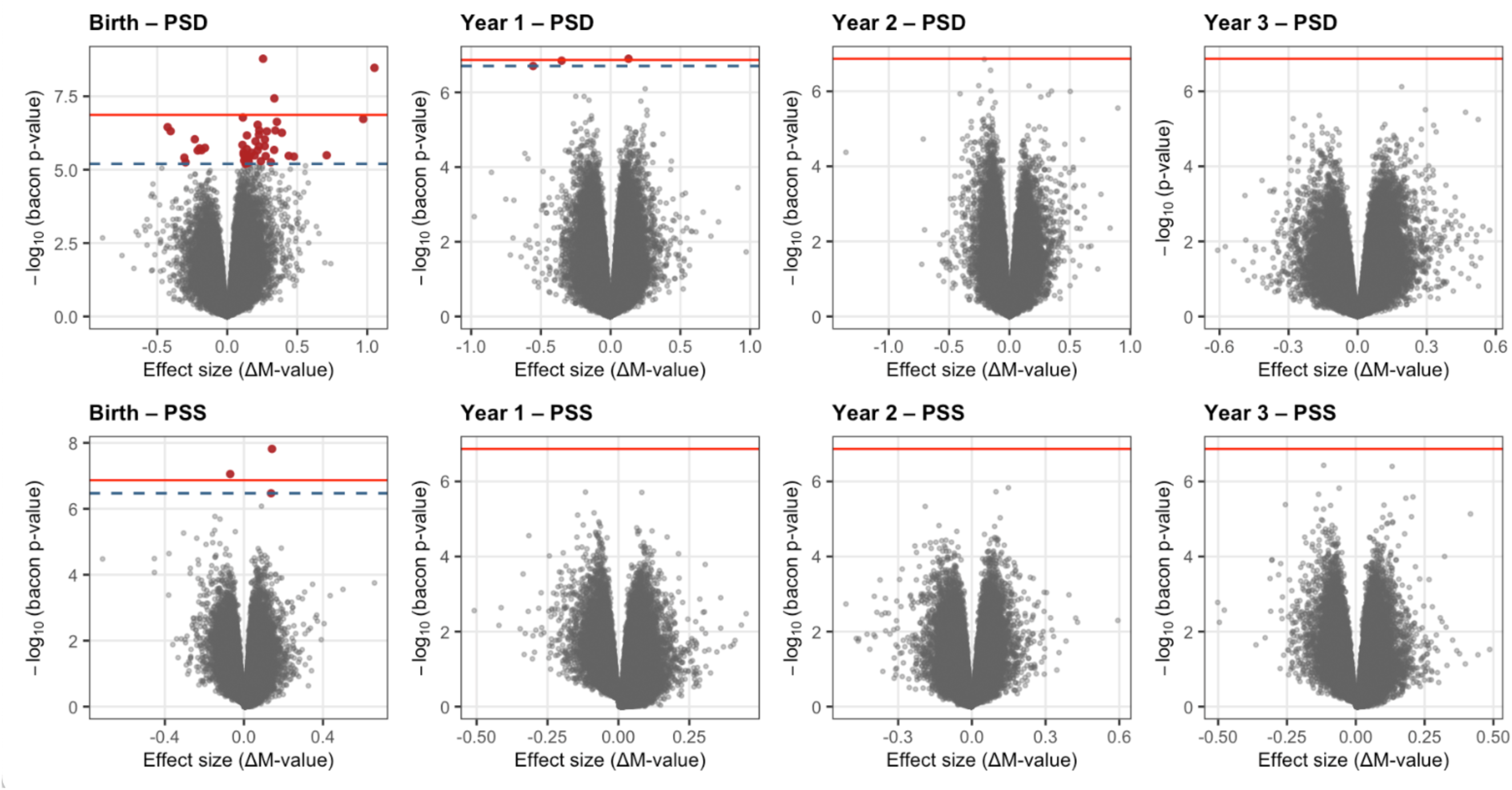
Genome-wide associations of prenatal exposures with child DNA methylation. Top row: Prenatal social disadvantage (PSD); Bottom row: Prenatal psychosocial stress (PSS); Red line = Bonferroni; Dashed blue line = FDR < 0.05; Red dots = FDR-significant CpGs; Bacon-corrected p-values shown for 7 panels; Hand-calculated FDR p-values shown for PSD Year 3 (see Methods)

#### Prenatal Psychosocial Stress

PSS was associated with three FDR-significant CpGs at birth: cg14800753 (β=0.14, bacon *p*=1.53 x 10^-08^), cg08019911 (β=0.14, bacon *p*=3.38 x 10^-07^), and cg12686847 (β=-0.07, bacon *p*=8.83 x 10^-08^). There were no significant CpGs at years 1, 2, or 3.

#### Persistence of Adversity-Associated Methylation Across Early Childhood

To examine whether PSD and PSS-associated methylation signatures persisted across early childhood, we identified all CpGs reaching Bacon-corrected FDR significance at birth or year 1 for PSD, along with the 3 significant CpGs at the birth timepoint for PSS and traced their Bacon-corrected p-values and effect sizes across all four timepoints. PSS-associated CpGs did not show any persistence across timepoints. In contrast, 13 PSD-associated CpGs maintained significance (*p*<0.05) across all four timepoints (**Table 2)**. An additional 14 CpGs (28%) showed nominal significance at three of four timepoints and 11 CpGs (22%) at two timepoints. Overall, 76% of discovery-set CpGs showed nominal significance at two or more timepoints, and direction of effect was preserved across all four timepoints with no instances of effect reversal. Among the 13 persistently associated CpGs, six were consistently hypomethylated (i.e., β consistently negative) in association with higher PSD and seven were consistently hypermethylated (**Figure 2**; **Figure 3**).

**Figure 2:**
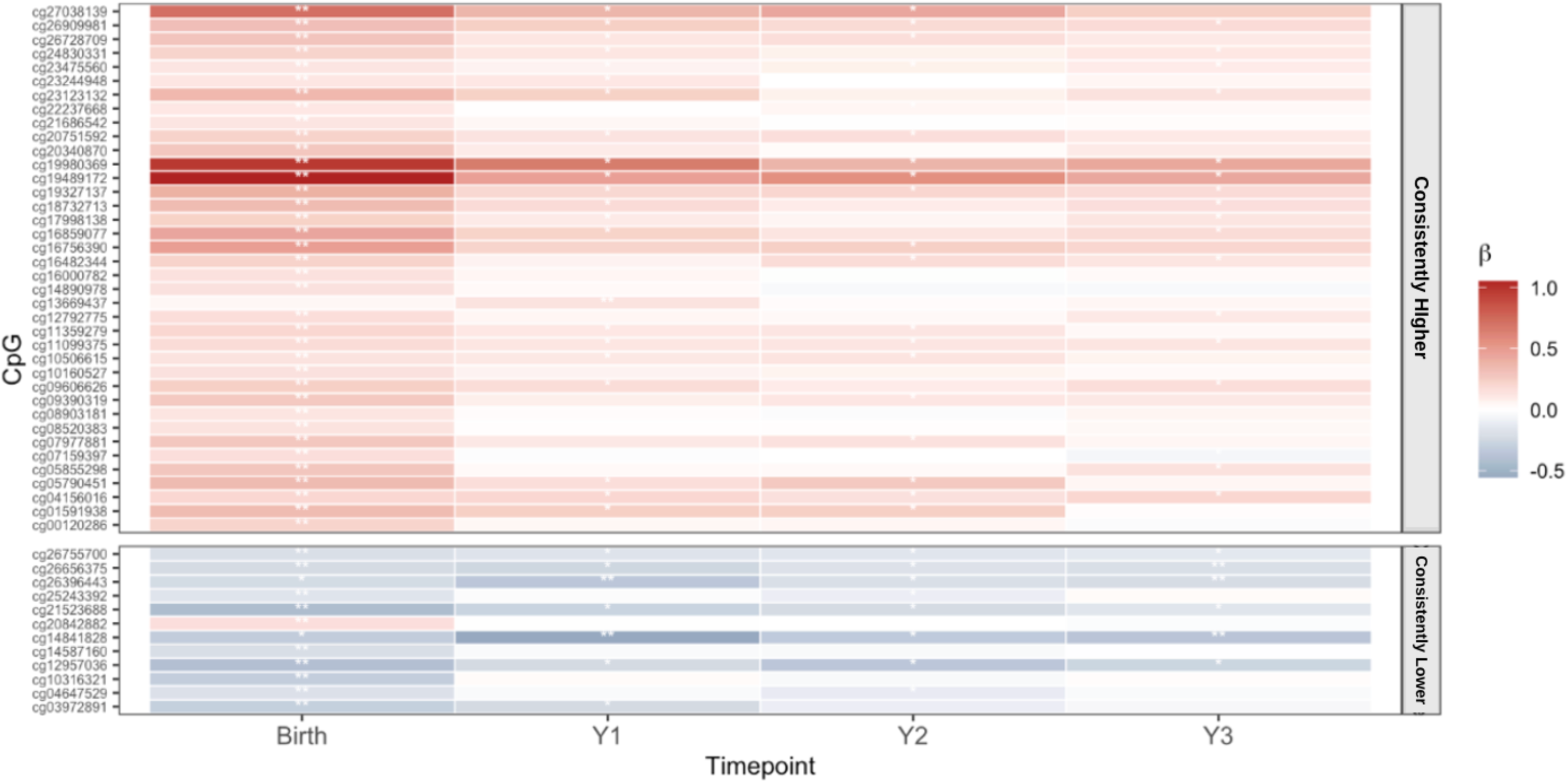
Epigenetic Effect Sizes Across Timepoints. ** Bacon FDR <0.05; * Bacon *p* < 0.05; CpGs color-coded by persistence and sorted by direction.

**Figure 3:**
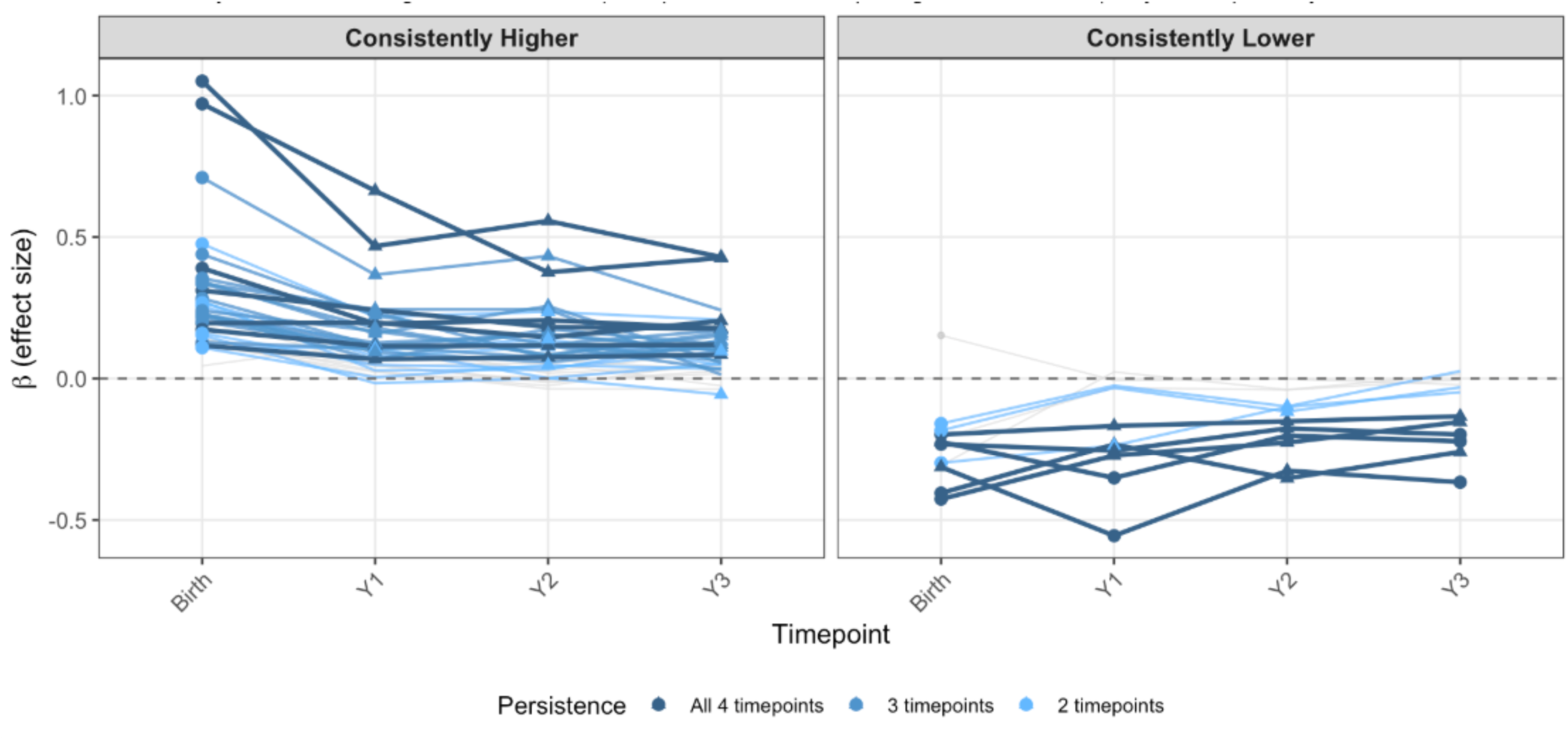
Beta Trajectories: Persistence of Epigenetic Signal. Color intensity = Bacon *p* < 0.05 significance across timepoints; Circle = Bacon FDR; Triangle = Bacon *p* < 0.05; Grey = 1 timepoint only.

**Table 2.**
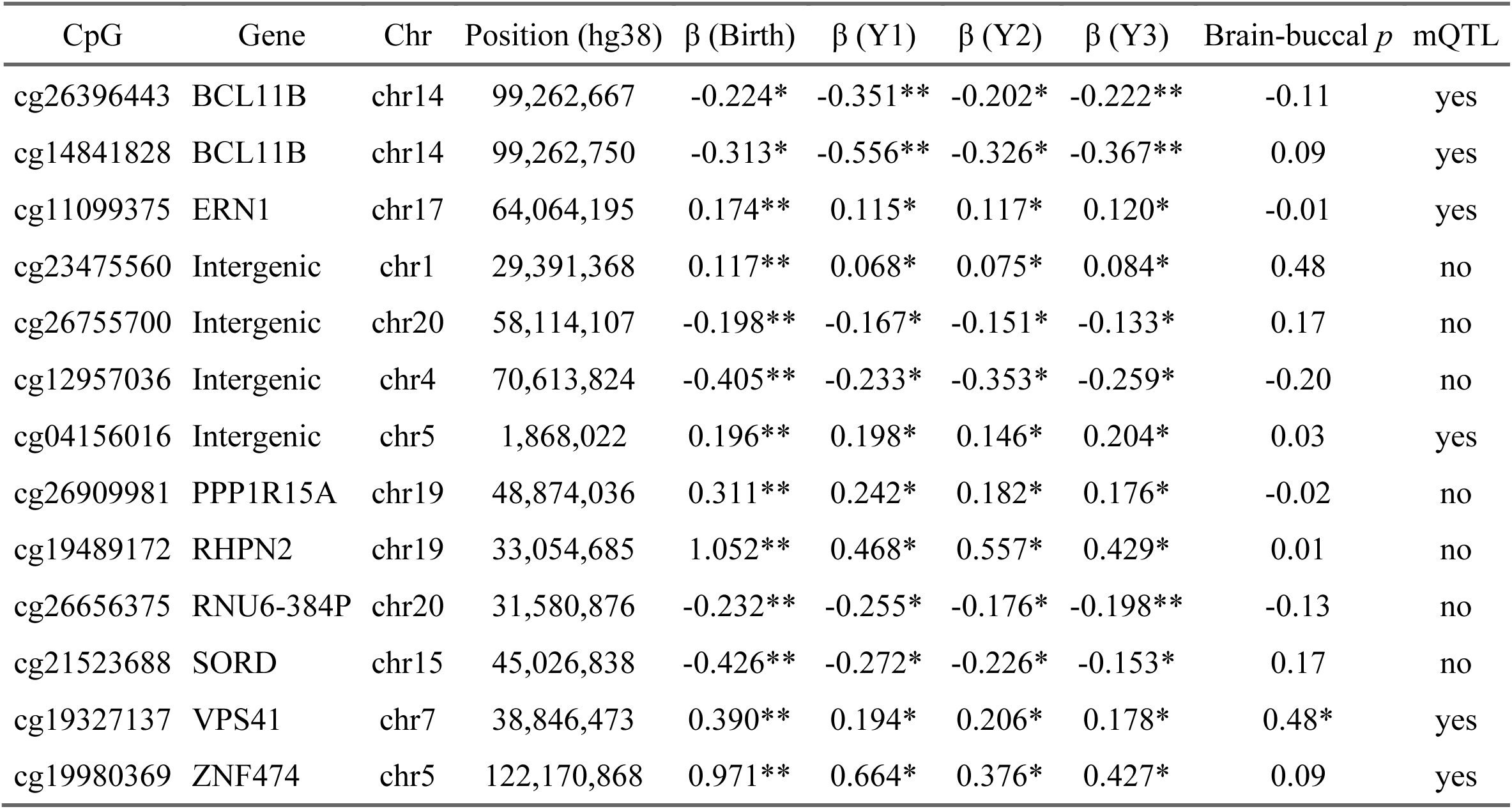
Thirteen CpGs Persistently associated (bacon-corrected p < 0.05) with Prenatal Social Disadvantage across all four timepoints. ** Bacon-corrected FDR < 0.05; * Bacon-corrected nominal p < 0.05. Effect sizes (β) on the M-value scale. All models adjusted for infant sex, infant race, maternal tobacco, CD45-positive cell proportion, maternal age at delivery, and gestational age at birth (birth timepoint) or child age at sample collection (postnatal timepoints). Brain–buccal ρ from IMAGE-CpG; * indicates *p* < 0.05 for tissue concordance. mQTL status from GoDMC.

| CpG | Gene | Chr | Position (hg38) | $\beta$ (Birth) | $\beta$ (Y1) | $\beta$ (Y2) | $\beta$ (Y3) | Brain-buccal $p$ | mQTL |
| --- | --- | --- | --- | --- | --- | --- | --- | --- | --- |
| cg26396443 | BCL11B | chr14 | 99,262,667 | -0.224* | -0.351** | -0.202* | -0.222** | -0.11 | yes |
| cg14841828 | BCL11B | chr14 | 99,262,750 | -0.313* | -0.556** | -0.326* | -0.367** | 0.09 | yes |
| cg11099375 | ERN1 | chr17 | 64,064,195 | 0.174** | 0.115* | 0.117* | 0.120* | -0.01 | yes |
| cg23475560 | Intergenic | chr1 | 29,391,368 | 0.117** | 0.068* | 0.075* | 0.084* | 0.48 | no |
| cg26755700 | Intergenic | chr20 | 58,114,107 | -0.198** | -0.167* | -0.151* | -0.133* | 0.17 | no |
| cg12957036 | Intergenic | chr4 | 70,613,824 | -0.405** | -0.233* | -0.353* | -0.259* | -0.20 | no |
| cg04156016 | Intergenic | chr5 | 1,868,022 | 0.196** | 0.198* | 0.146* | 0.204* | 0.03 | yes |
| cg26909981 | PPP1R15A | chr19 | 48,874,036 | 0.311** | 0.242* | 0.182* | 0.176* | -0.02 | no |
| cg19489172 | RHPN2 | chr19 | 33,054,685 | 1.052** | 0.468* | 0.557* | 0.429* | 0.01 | no |
| cg26656375 | RNU6-384P | chr20 | 31,580,876 | -0.232** | -0.255* | -0.176* | -0.198** | -0.13 | no |
| cg21523688 | SORD | chr15 | 45,026,838 | -0.426** | -0.272* | -0.226* | -0.153* | 0.17 | no |
| cg19327137 | VPS41 | chr7 | 38,846,473 | 0.390** | 0.194* | 0.206* | 0.178* | 0.48* | yes |
| cg19980369 | ZNF474 | chr5 | 122,170,868 | 0.971** | 0.664* | 0.376* | 0.427* | 0.09 | yes |

### Functional Enrichment of Epigenetic Signatures

We performed functional enrichment, a way of characterizing the genes and proteins associated with a specific CpG site, of all 50 FDR-significant CpGs from the EWAS using Enrichr^29^.

**Figure 4** summarizes the gene annotation and relevant biological pathways associated with the 50 CpGs (30 unique protein-coding genes, after excluding pseudogenes and non-coding annotations). Regarding biological processes (e.g., WikiPathways), three pathways survived FDR correction: unfolded protein response (UPR), photodynamic therapy-induced unfolded protein response, and endoplasmic reticulum (ER) stress response in coronavirus infection (all p_adj_ < 0.024). Each path was driven by the same two genes, *ERN1* and *PPP1R15A*, suggesting a convergent signal. The same gene pair produced the strongest Gene Ontology (GO) term, intrinsic apoptotic signaling in response to ER stress, which reached nominal but not FDR- adjusted significance.

**Figure 4:**
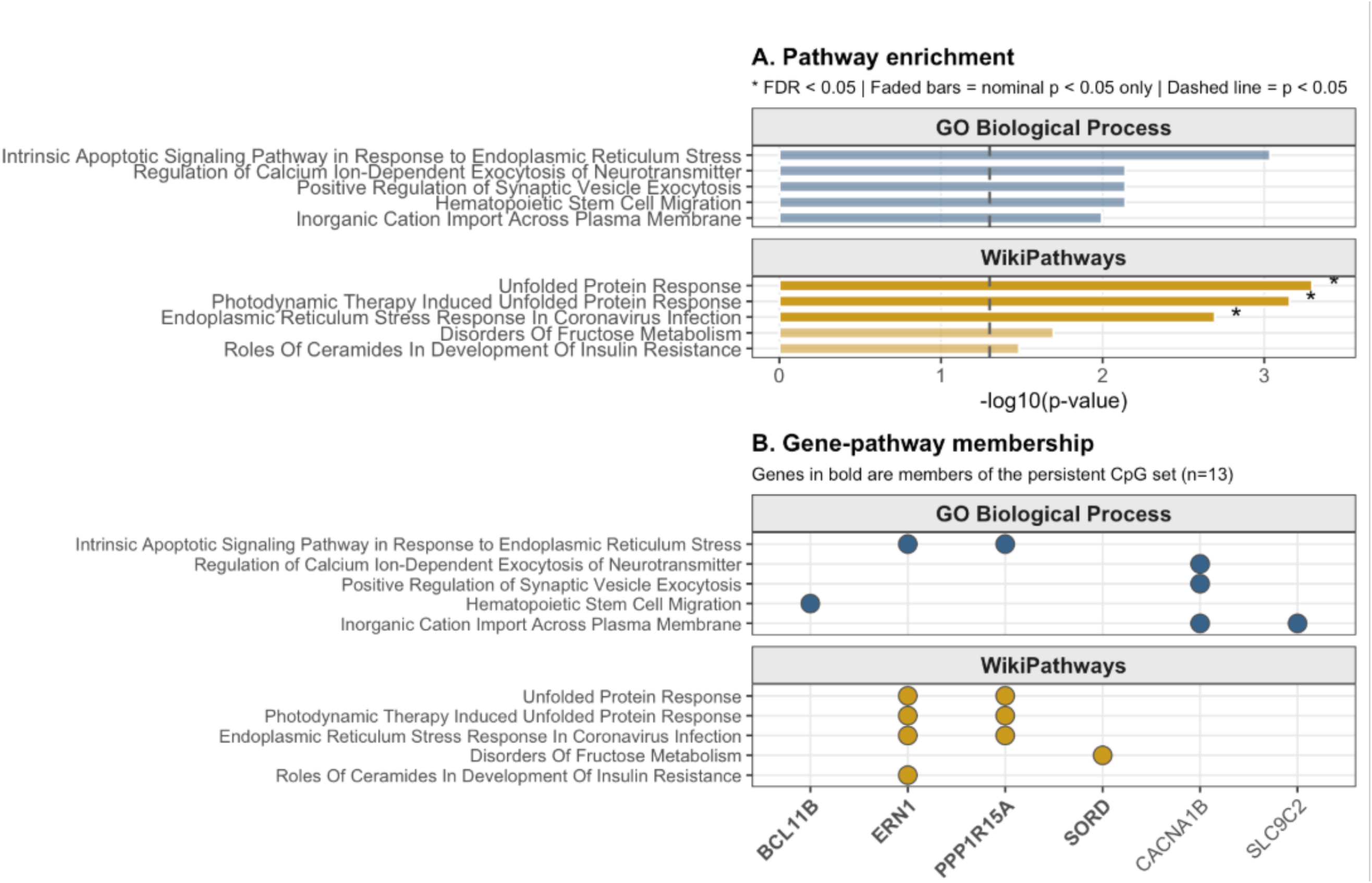
Enrichr Results.

### Tissue Type and Genetic Confounding

Given that saliva is a peripheral tissue, we assessed the extent to which methylation at the 13 persistent CpG sites was influenced by underlying heritability (e.g., genotype). To that end, we cross-referenced each CpG against GoDMC^34^, a large blood-based catalogue of methylation quantitative trait loci (mQTL), which indicate that a given CpG may be driven primarily by genetic rather than environmental influence. Six of the 13 persistent CpGs had documented mQTLs, including both *BCL11B* promoter sites (cg26396443, cg14841828), *ERN1* (cg11099375), and *ZNF474* (cg19980369).

### Mediation of Prenatal Disadvantage Effects via Birth Methylation

We examined whether DNA methylation at birth mediated associations between prenatal exposures and child psychopathology at ages four-six on a subset of children with available phenotype data (n=140). We conducted mediation analyses using a pre-specified subset of candidate CpGs: the thirteen ‘persistent’ CpGs identified above and the PSS-associated FDR-significant CpGs at birth. ‘Persistent’ set CpGs were tested with PSD as the exposure; PSS-associated CpGs were tested with PSS as the exposure. In adjusted models (covariates: child sex, race, gestational age at birth), methylation at cg19980369 (*ZNF474*) showed an uncorrected negative indirect effect of PSD on externalizing symptoms (ACME = −0.76, *p* = 0.028).

However, this association was not significant following FDR correction and therefor results should be considered exploratory. Complete mediation results are provided in **Table 3**.

**Table 3:** Mediation of prenatal exposure effects on child psychopathology via birth DNA methylation. ACME = average causal mediation effect. 95% CIs from 1,000 nonparametric bootstrap iterations. Unadjusted models include only exposure → mediator → outcome. Adjusted models additionally control for child sex, race, and gestational age at birth. *p* = bootstrap *p*-value for the ACME. *q* = Benjamini-Hochberg FDR within outcome family. PSD = prenatal social disadvantage; PSS = prenatal psychosocial stress. Persistent PSD-associated CpGs tested with PSD as exposure; PSS birth-associated CpGs tested with PSS as exposure. Outcomes: PAPA symptom counts at year 4-6. N = 140 children with both birth methylation and year 4-6 PAPA data

| Exposure | CpG (gene) | Outcome | N | ACME (unadj.) [95% CI] | p (unadj.) | q (unadj.) | ACME (adj.) [95% CI] | p (adj.) | q (adj.) |
| --- | --- | --- | --- | --- | --- | --- | --- | --- | --- |
| PSD | cg19980369 (ZNF474) | Externalizing | 140 | -0.980 [-1.85, -0.13] | 0.018 | 0.270 | -0.761 [-1.48, -0.07] | 0.028 | 0.360 |
| PSD | cg04156016 ((Intergenic)) | Externalizing | 140 | -0.443 [-0.99, -0.01] | 0.048 | 0.320 | -0.509 [-1.14, 0.04] | 0.072 | 0.360 |
| PSD | cg23475560 ((Intergenic)) | Externalizing | 140 | -0.443 [-1.01, 0.01] | 0.064 | 0.320 | -0.412 [-1.03, 0.04] | 0.072 | 0.360 |
| PSD | cg26656375 (RNU6-384P) | Externalizing | 140 | -0.384 [-1.27, 0.52] | 0.376 | 0.627 | -0.465 [-1.17, 0.25] | 0.194 | 0.636 |
| PSD | cg19489172 (RHPN2) | Externalizing | 140 | -0.677 [-1.78, 0.44] | 0.244 | 0.604 | -0.597 [-1.68, 0.35] | 0.212 | 0.636 |
| PSD | cg26755700 ((Intergenic)) | Externalizing | 140 | -0.439 [-1.20, 0.30] | 0.258 | 0.604 | -0.335 [-0.91, 0.24] | 0.268 | 0.670 |
| PSD | cg12957036 ((Intergenic)) | Externalizing | 140 | -0.495 [-1.29, 0.30] | 0.250 | 0.604 | -0.318 [-0.94, 0.37] | 0.370 | 0.793 |
| PSD | cg26909981 (PPP1R15A) | Externalizing | 140 | 0.598 [-0.44, 1.64] | 0.282 | 0.604 | 0.339 [-0.53, 1.23] | 0.450 | 0.844 |
| PSD | cg21523688 (SORD) | Externalizing | 140 | -0.243 [-1.16, 0.60] | 0.546 | 0.705 | -0.210 [-0.95, 0.47] | 0.532 | 0.852 |
| PSD | cg26396443 (BCL11B) | Externalizing | 140 | 0.093 [-0.31, 0.47] | 0.682 | 0.787 | 0.066 [-0.21, 0.34] | 0.602 | 0.852 |
| PSD | cg19327137 (VPS41) | Externalizing | 140 | -0.277 [-1.35, 0.63] | 0.564 | 0.705 | -0.191 [-1.21, 0.78] | 0.738 | 0.852 |
| PSD | cg14841828 (BCL11B) | Externalizing | 140 | 0.017 [-0.24, 0.23] | 0.894 | 0.958 | 0.018 [-0.18, 0.16] | 0.880 | 0.888 |
| PSD | cg11099375 (ERN1) | Externalizing | 140 | -0.008 [-0.71, 0.61] | 0.998 | 0.998 | 0.030 [-0.51, 0.55] | 0.888 | 0.888 |
| PSD | cg21523688 (SORD) | Internalizing | 140 | -0.281 [-0.76, 0.14] | 0.226 | 0.866 | -0.255 [-0.68, 0.10] | 0.178 | 0.945 |
| PSD | cg26656375 (RNU6-384P) | Internalizing | 140 | -0.172 [-0.66, 0.29] | 0.462 | 0.866 | -0.230 [-0.63, 0.14] | 0.232 | 0.945 |
| PSD | cg19980369 (ZNF474) | Internalizing | 140 | -0.211 [-0.64, 0.20] | 0.326 | 0.866 | -0.178 [-0.50, 0.14] | 0.268 | 0.945 |
| PSD | cg19489172 (RHPN2) | Internalizing | 140 | -0.238 [-0.77, 0.31] | 0.398 | 0.866 | -0.233 [-0.71, 0.23] | 0.330 | 0.945 |
| PSD | cg12957036 ((Intergenic)) | Internalizing | 140 | -0.240 [-0.69, 0.23] | 0.336 | 0.866 | -0.174 [-0.55, 0.20] | 0.364 | 0.945 |
| PSD | cg19327137 (VPS41) | Internalizing | 140 | -0.245 [-0.69, 0.17] | 0.268 | 0.866 | -0.209 [-0.65, 0.22] | 0.378 | 0.945 |
| PSD | cg04156016 ((Intergenic)) | Internalizing | 140 | -0.082 [-0.32, 0.13] | 0.430 | 0.866 | -0.073 [-0.35, 0.20] | 0.586 | 0.954 |
| PSD | cg26755700 ((Intergenic)) | Internalizing | 140 | -0.072 [-0.42, 0.31] | 0.724 | 0.980 | -0.068 [-0.33, 0.25] | 0.636 | 0.954 |
| PSD | cg14841828 (BCL11B) | Internalizing | 140 | -0.048 [-0.19, 0.07] | 0.456 | 0.866 | -0.024 [-0.13, 0.05] | 0.640 | 0.954 |
| PSD | cg26909981 (PPP1R15A) | Internalizing | 140 | 0.009 [-0.48, 0.49] | 0.980 | 0.980 | -0.085 [-0.48, 0.32] | 0.712 | 0.954 |
| PSD | cg23475560 ((Intergenic)) | Internalizing | 140 | 0.016 [-0.21, 0.25] | 0.900 | 0.980 | 0.011 [-0.20, 0.24] | 0.924 | 0.954 |
| PSD | cg26396443 (BCL11B) | Internalizing | 140 | 0.003 [-0.21, 0.21] | 0.976 | 0.980 | -0.002 [-0.15, 0.15] | 0.946 | 0.954 |
| PSD | cg11099375 (ERN1) | Internalizing | 140 | -0.020 [-0.37, 0.32] | 0.862 | 0.980 | -0.008 [-0.30, 0.28] | 0.954 | 0.954 |
| PSS | cg12686847 (HSPA1L) | Externalizing | 140 | -0.150 [-0.50, 0.17] | 0.346 | 0.627 | -0.091 [-0.42, 0.26] | 0.632 | 0.852 |
| PSS | cg14800753 (EXOC4) | Externalizing | 140 | -0.118 [-0.60, 0.26] | 0.546 | 0.705 | -0.076 [-0.57, 0.38] | 0.724 | 0.852 |
| PSS | cg14800753 (EXOC4) | Internalizing | 140 | -0.015 [-0.18, 0.13] | 0.800 | 0.980 | 0.024 [-0.16, 0.22] | 0.794 | 0.954 |
| PSS | cg12686847 (HSPA1L) | Internalizing | 140 | -0.023 [-0.17, 0.18] | 0.792 | 0.980 | 0.008 [-0.16, 0.21] | 0.920 | 0.954 |

## Discussion

We characterized the epigenetic signatures of two forms of prenatal adversity, prenatal social disadvantage (PSD) and prenatal psychosocial stress (PSS), across four cross-sectional timepoints spanning birth through age three in the eLABE cohort. We further tested any mediating effects of significant CpGs in the pathway from prenatal exposures to child internalizing and externalizing symptoms at 4-6 years old.

We present three findings extending the literature on the associations among prenatal adversity, child DNA methylation, and early childhood psychopathology. First, both PSD and PSS were associated with epigenetic signatures in offspring. Second, 13 CpGs associated with PSD at birth and year 1 retained significance across all four timepoints from birth to three, whereas PSS-associated CpGs did not show a persistent effect across time. The persistently-associated PSD CpGs showed an association with cellular stress response biology, most notably the unfolded protein response (UPR), driven by CpGs mapping to *ERN1* and *PPP1R15A*. Third, exploratory mediation analyses showed an uncorrected indirect effect of methylation at *ZNF474* (cg19980369) on the association between PSD and externalizing symptoms at ages 4-6, although this finding did not survive multiple-testing correction. In summary, PSD-associated marks are detectable at birth, persist into early childhood, converge on biologically coherent stress-response pathways. While we did not find significant mediation effects in our sample, future work should leverage larger developmental cohorts to interrogate this mediating pathway with greater statistical power.

### Social Disadvantage, Stress, and the Epigenome

PSD and PSS demonstrated divergent offspring DNAm profiles. This difference is striking given the robust literature on the associations between prenatal stress and offspring psychopathology. However, structural disadvantage, which tends to show little fluctuation across prenatal and postpartum time points (without intervention) may pose a more consequential risk to pre- and postnatal development, with lingering effects into the early years of life. The effects of maternal stress, which may be more episodic or phasic in nature compared with structural disadvantage, may also be more easily tempered with positive postnatal experiences such as secure attachment and positive parenting. The persistent effects of PSD on DNAm aligns with the previously proposed concept of a biological poverty line: a threshold below which the developing brain is shaped by experience along qualitatively distinct trajectories and may act as persistent recalibration of the developmental context^35^. The convergence of the persistently-associated PSD CpGs on cellular stress-response biology is congruent with prior evidence that disadvantage-related inflammatory signaling is associated with neonatal brain microstructure in this cohort^36^, pointing to sustained physiological pathways through which structural adversity may be embedded. Ongoing research is testing the ‘essential ingredients’ necessary to buffer the impact of social disadvantage and preserve healthy fetal and infant neurodevelopment. For instance, the thrive factor (T-factor), encompassing safety, sensitive caregiving, healthy sleep, good nutrition, and environmental stimulation, has shown positive associations with early childhood cognition even when accounting for the effects of social disadvantage^37^. The promotive effects of the T-factor were themselves stratified by disadvantage, such that the cognitive benefits associated with the T-factor were more evident for children at low and average levels of PSD, but not for children facing the highest levels of PSD – suggesting that a baseline level of environmental support may be necessary before positive inputs can be tangibly beneficial. Our findings echo this framework at the molecular level in our observation that PSD, but not PSS, leaves a persistent DNAm signature from birth through year 3.

### Prenatal Adversity and Offspring DNAm

Our finding that PSD is associated with epigenetic signatures at birth and year 1 is consistent with a growing body of research linking prenatal adversity to offspring DNAm. Prior literature has found associations between prenatal socioeconomic disadvantage^38 39^ as well as maternal education^40^ and offspring birth methylation. A recent meta-analysis of the international Pregnancy and Childhood Epigenetics consortium (PACE; *N*=5,496) found a convergent offspring epigenetic signal associated with a closely-related construct, maternal stressful life events during pregnancy^21^.

Our results add to this literature in two important ways. First, we found differences between structural versus psychological dimensions of prenatal adversity, with PSD producing over 15 times more FDR-significant CpGs at birth than PSS. One interpretation is that structural adversity captures a more chronic, biologically pervasive exposure than time-limited psychological stress; another is that experiential dimensions may exert effects through more localized pathways (e.g., HPA axis regulation) that are less detectable at the epigenome-wide level. Second, our multi-timepoint design during infancy and early childhood allows us to deeply characterize patterns of methylation during early neuroplastic periods. Most prior EWAS of prenatal adversity have examined a single timepoint, leaving unresolved whether observed associations represent stable or transient methylation responses that dissipate over time. Recent longitudinal work in the ALSPAC cohort has begun to address this gap: childhood adversity has been shown to produce time-dependent effects on early childhood DNAm^22 41^. This persistent effect may reflect enduring epigenetic programming established during fetal development, consistent with prior empirical work^42^ as well as the developmental origins of health and disease (DOHaD) framework^43^. Persistent effects may also reflect the ongoing nature of postnatal social disadvantage in children born to disadvantaged families. In other words, continued exposure to the same structural conditions that shaped their prenatal environment may contribute to persistence of a methylation signal into early childhood. Notably, we observed effect-size attenuation across time despite maintained direction and significance. This may be indicative of a gradual dilution of the prenatal signal or an artifact of reduced sample sizes at subsequent time points. The persistent epigenetic signature of PSD across the first three years of life reinforces the potential of DNAm as a candidate biomarker for tracking the biological embedding of early-life adversity, with implications for identifying developmental windows during which intervention may be most consequential.

### Gene annotation findings and neurodevelopmental relevance

The functional enrichment of PSD-associated CpGs at genes involved in the unfolded protein response (UPR) and endoplasmic reticulum (ER) stress signaling is a novel finding that has neurodevelopmental relevance. *ERN1* (IRE1α) is one of three canonical ER stress sensors and coordinates the adaptive cellular response to accumulated unfolded proteins^44 45^. *PPP1R15A* (GADD34) is a key effector of the integrated stress response that regulates translational recovery following cellular stress^46 47^. Both genes have been implicated in downstream neuronal function and psychiatric disease. Prolonged UPR activation contributes to neuronal apoptosis and has been linked to neurodegenerative disease^48^ and to schizophrenia and mood disorders^49^.

Preclinical work has demonstrated that early-life stress in rodents alters *Ern1* expression in the medial prefrontal cortex in early life^50^. The convergence of persistently disadvantage-associated CpGs on this pathway suggests that PSD may leave a molecular signature reflecting altered cellular stress-response pathways which may hold consequences for downstream neurodevelopmental maturation trajectories. The persistent PSD-associated CpGs at *BCL11B*, whose loss-of-function mutations have been associated with neurodevelopmental disability and autism^51,52^, further reinforce the neurodevelopmental relevance of the identified signature.

Together, this enrichment pattern is suggestive of a model in which PSD may become biologically embedded through pathways governing cellular stress response and neural developmental programming, though direct evidence linking these methylation changes to brain outcomes require complementary tissue-level and expression-level validation.

### Prenatal Adversity, Child DNAm, and Emerging Psychopathology

Our mediation analyses did not reveal any significant effects of child DNAm on the relationship between PSD and child psychopathology at ages 4-6. We did find one nominal indirect effect of methylation at *ZNF474* (cg19980369) on the association between PSD and year 4-6 externalizing symptoms. Several features are notable about cg19980369 which may make this exploratory finding relevant for future studies. It is one of the 13 CpGs that retained significance across all timepoints from birth to year 3, carried one of the largest effect sizes in the birth EWAS, and is associated with an mQTL. Our mediation models were likely underpowered; however, our exploratory finding was directionally consistent with patterns in other empirical work which have documented indirect effects of DNAm in the opposite direction of the positive associations with adversity and externalizing symptoms later in development, suggesting a suppression effect^22,53^. Replication in larger samples is needed. Larger cohorts with harmonized methylation and outcome measures, such as those that will likely be available through initiatives like the Pregnancy and Childhood Epigenetics (PACE)^54^ Consortium, the HEALthy Brain and Child Development Study (HBCD)^55^, and the Methylation, Imaging, and NeuroDevelopment (MIND)^56^ Consortium, will be well-positioned to test these mediation hypotheses at scale.

### Strengths and Limitations

This study has several strengths. First, this cohort affords a unique opportunity to characterize the entire transmission pathway from pregnancy through early childhood. Further, this is the first study to our knowledge to capture serial DNAm measures during the window from birth through preschool age. Second, our within-cohort comparison of two conceptually related but distinct dimensions of prenatal adversity (PSD versus PSS) provides a within-sample test of biological specificity that has been largely absent from prior work. Third, our cohort was explicitly oversampled for PSD, resulting in approximately half of the mother-infants in our study being affected by poverty and thereby allowing us to isolate the effects of environmental adversity on biological outcomes.

Several limitations warrant consideration. First, our sample sizes at each timepoint (n=130-215) are modest by EWAS standards, although we employed a Bayesian approach to boost statistical power. Second, saliva is a peripheral tissue and should therefore be interpreted carefully, although some significant CpGs were identified to have brain-buccal concordance using one of the only saliva-brain reference databases available. Third, we could not resolve the anomalous bacon bias parameter at the year 3 PSD model, which may reflect latent batch effects or unmodeled confounders at that timepoint. Finally, we cannot fully account for genetic ancestry (given we did not include genetic principal components into the current analysis) or postnatal environmental exposures, and the potential contribution of methylation quantitative trait loci (mQTLs) to observed associations warrants further investigation as forthcoming genotype data become available. Future analyses (Mendelian randomization; examining pre- vs postnatal influences) could help address these limitations.

## Conclusion

PSD was associated with a robust, biologically coherent, and developmentally persistent DNA methylation signature in early childhood, with convergent involvement of the UPR, ER stress signaling, and cortical neurodevelopmental transcription factors. These findings advance a mechanistic understanding of how structural dimensions of prenatal adversity may become biologically embedded during early neurodevelopment and suggest that DNA methylation may function as one component of a broader biological pathway linking prenatal environment to child mental health outcomes. Future work integrating methylation, gene expression, brain imaging, and multi-timepoint psychopathology in larger cohorts will be essential to move from association to mechanism, and ultimately to inform the timing and targeting of preventive interventions during the earliest windows of neurodevelopment.

## Notes

### Competing Interest Statement

The authors have declared no competing interest.

### Funding

- National Institute of Mental Health: 5R01MH113883-09, 5T32MH100019-08

## References

1. Crear-Perry J, Correa-de-Araujo R, Lewis Johnson T, McLemore MR, Neilson E, Wallace M. Social and Structural Determinants of Health Inequities in Maternal Health. J Womens Health. 2021;30(2):230–235. doi:10.1089/jwh.2020.8882

2. Herzberg MP, Smyser CD. Prenatal Social Determinants of Health: Narrative review of maternal environments and neonatal brain development. Pediatr Res. 2024;96(6):1417–1428. doi:10.1038/s41390-024-03345-7

3. Lean RE, Constantino-Pettit A, Gorham LS, et al. Social Determinants of Health, the developing brain, and risk and resilience for psychopathology. Neuropsychopharmacology. 2026;51(1):185–202. doi:10.1038/s41386-025-02169-1

4. Gilman SE, Hornig M, Ghassabian A, et al. Socioeconomic disadvantage, gestational immune activity, and neurodevelopment in early childhood. Proc Natl Acad Sci. 2017;114(26):6728–6733. doi:10.1073/pnas.1617698114

5. Glynn LM, Sandman CA. Prenatal Origins of Neurological Development: A Critical Period for Fetus and Mother. Curr Dir Psychol Sci. 2011;20(6):384–389. doi:10.1177/0963721411422056

6. Gilmore JH, Knickmeyer RC, Gao W. Imaging structural and functional brain development in early childhood. Nat Rev Neurosci. 2018;19(3):123–137. doi:10.1038/nrn.2018.1

7. Triplett RL, Lean RE, Parikh A, et al. Prenatal Exposure to Early-Life Adversity and Neonatal Brain Volumes at Birth. Accessed June 25, 2026. https://jamanetwork.com/journals/jamanetworkopen/fullarticle/2790989

8. Koyama Y, Hidalgo APC, Lacey RE, et al. Poverty from fetal life onward and child brain morphology. Sci Rep. 2023;13(1):1295. doi:10.1038/s41598-023-28120-2

9. Johnson SB, Riis JL, Noble KG. State of the Art Review: Poverty and the Developing Brain. Pediatrics. 2016;137(4):e20153075. doi:10.1542/peds.2015-3075

10. Cook KM, De Asis-Cruz J, Kapse K, et al. Prenatal experience of greater neighborhood disadvantage is associated with altered fetal volumetric brain growth in utero. Cereb Cortex. 2026;36(3):bhag017. doi:10.1093/cercor/bhag017

11. Lean RE, Smyser CD, Brady RG, et al. Prenatal exposure to maternal social disadvantage and psychosocial stress and neonatal white matter connectivity at birth. Proc Natl Acad Sci. 2022;119(42):e2204135119. doi:10.1073/pnas.2204135119

12. Demers CH, Bagonis MM, Al-Ali K, et al. Exposure to prenatal maternal distress and infant white matter neurodevelopment. Dev Psychopathol. 2021;33(5):1526–1538. doi:10.1017/S0954579421000742

13. Nielsen AN, Triplett RL, Bernardez LM, et al. Prenatal social disadvantage is associated with alterations in functional networks at birth. Proc Natl Acad Sci. 2024;121(50):e2405448121. doi:10.1073/pnas.2405448121

14. Leverett SD, Brady RG, Tooley UA, et al. Associations between Parenting and Cognitive and Language Abilities at 2 Years of Age Depend on Prenatal Exposure to Disadvantage. J Pediatr. 2025;276:114289. doi:10.1016/j.jpeds.2024.114289

15. Nelson CA, Gabard-Durnam LJ. Early Adversity and Critical Periods: Neurodevelopmental Consequences of Violating the Expectable Environment. Trends Neurosci. 2020;43(3):133–143. doi:10.1016/j.tins.2020.01.002

16. Monk C, Lugo-Candelas C, Trumpff C. Prenatal Developmental Origins of Future Psychopathology: Mechanisms and Pathways. Annu Rev Clin Psychol. 2019;15(1):317–344. doi:10.1146/annurev-clinpsy-050718-095539

17. Merrill SM, Hogan C, Bozack AK, et al. Telehealth Parenting Program and Salivary Epigenetic Biomarkers in Preschool Children With Developmental Delay: NIMHD Social Epigenomics Program. JAMA Netw Open. 2024;7(7):e2424815. doi:10.1001/jamanetworkopen.2024.24815

18. Weaver IC g., Diorio J, Seckl JR, Szyf M, Meaney MJ. Early Environmental Regulation of Hippocampal Glucocorticoid Receptor Gene Expression: Characterization of Intracellular Mediators and Potential Genomic Target Sites. Ann N Y Acad Sci. 2004;1024(1):182–212. doi:10.1196/annals.1321.099

19. Liff CW, Ayman YR, Jaeger EC, et al. Fear conditioning biases olfactory sensory neuron frequencies across generations. Penzo MA, Wassum KM, eds. eLife. 2026;12:RP92882. doi:10.7554/eLife.92882

20. Segura AG, de la Serna E, Sugranyes G, et al. Epigenetic age deacceleration in youth at familial risk for schizophrenia and bipolar disorder. Transl Psychiatry. 2023;13(1):155. doi:10.1038/s41398-023-02463-w

21. Kotsakis Ruehlmann A, Sammallahti S, Cortés Hidalgo AP, et al. Epigenome-wide meta-analysis of prenatal maternal stressful life events and newborn DNA methylation. Mol Psychiatry. 2023;28(12):5090–5100. doi:10.1038/s41380-023-02010-5

22. Lussier AA, Zhu Y, Smith BJ, et al. Association between the timing of childhood adversity and epigenetic patterns across childhood and adolescence: findings from the Avon Longitudinal Study of Parents and Children (ALSPAC) prospective cohort. Lancet Child Adolesc Health. 2023;7(8):532–543. doi:10.1016/S2352-4642(23)00127-X

23. Abrishamcar S, Zhuang BC, Thomas M, et al. Association between maternal perinatal stress and depression and infant DNA methylation in the first year of life. Transl Psychiatry. 2024;14(1):445. doi:10.1038/s41398-024-03148-8

24. Barker ED, Walton E, Cecil CAM. Annual Research Review: DNA methylation as a mediator in the association between risk exposure and child and adolescent psychopathology. J Child Psychol Psychiatry. 2018;59(4):303–322. doi:10.1111/jcpp.12782

25. Luby JL, England SK, Barch DM, et al. Social disadvantage during pregnancy: effects on gestational age and birthweight. J Perinatol. 2023;43(4):477–483. doi:10.1038/s41372-023-01643-2

26. Min JL, Hemani G, Davey Smith G, Relton C, Suderman M. Meffil: efficient normalization and analysis of very large DNA methylation datasets. Bioinformatics. 2018;34(23):3983–3989. doi:10.1093/bioinformatics/bty476

27. DelCarmen-Wiggins R, Carter A. Handbook of Infant, Toddler, and Preschool Mental Health Assessment. Oxford University Press; 2004.

28. Du P, Zhang X, Huang CC, et al. Comparison of Beta-value and M-value methods for quantifying methylation levels by microarray analysis. BMC Bioinformatics. 2010;11(1):587. doi:10.1186/1471-2105-11-587

29. Kuleshov MV, Jones MR, Rouillard AD, et al. Enrichr: a comprehensive gene set enrichment analysis web server 2016 update. Nucleic Acids Res. 2016;44(W1):W90–W97. doi:10.1093/nar/gkw377

30. Braun P, Han S, Nagahama Y, et al. 28 - IMAGE-CpG: DEVELOPMENT OF A WEB-BASED SEARCH TOOL FOR GENOME-WIDE DNA METHYLATION CORRELATION BETWEEN LIVE HUMAN BRAIN AND PERIPHERAL TISSUES WITHIN INDIVIDUALS. Eur Neuropsychopharmacol. 2019;29:S796. doi:10.1016/j.euroneuro.2017.08.029

31. van Iterson M, van Zwet EW, Heijmans BT, the BIOS Consortium. Controlling bias and inflation in epigenome- and transcriptome-wide association studies using the empirical null distribution. Genome Biol. 2017;18(1):19. doi:10.1186/s13059-016-1131-9

32. Barfield RT, Kilaru V, Smith AK, Conneely KN. CpGassoc: an R function for analysis of DNA methylation microarray data. Bioinformatics. 2012;28(9):1280–1281. doi:10.1093/bioinformatics/bts124

33. Tingley D, Yamamoto T, Hirose K, Keele L, Imai K. mediation: R Package for Causal Mediation Analysis. J Stat Softw. 2014;059(i05). doi:10.18637/jss.v059.i05

34. Min JL, Hemani G, Hannon E, et al. Genomic and phenotypic insights from an atlas of genetic effects on DNA methylation. Nat Genet. 2021;53(9):1311–1321. doi:10.1038/s41588-021-00923-x

35. Luby JL, Rank MR, Barch DM. Biological Poverty Line for Infants—Evidence and Implications. JAMA Pediatr. 2024;178(6):516–517. doi:10.1001/jamapediatrics.2024.0651

36. Sanders AFP, Tirado B, Seider NA, et al. Prenatal exposure to maternal disadvantage-related inflammatory biomarkers: associations with neonatal white matter microstructure. Transl Psychiatry. 2024;14(1):72. doi:10.1038/s41398-024-02782-6

37. Luby JL, Herzberg MP, Hoyniak C, et al. Basic Environmental Supports for Positive Brain and Cognitive Development in the First Year of Life. JAMA Pediatr. 2024;178(5):465–472. doi:10.1001/jamapediatrics.2024.0143

38. Simanek AM, Manansala R, Woo JMP, Meier HCS, Needham BL, Auer PL. Prenatal Socioeconomic Disadvantage and Epigenetic Alterations at Birth Among Children Born to White British and Pakistani Mothers in the Born in Bradford Study. Epigenetics. 2022;17(13):1976–1990. doi:10.1080/15592294.2022.2098569

39. Laubach ZM, Perng W, Cardenas A, et al. Socioeconomic Status and DNA Methylation from Birth Through Mid-Childhood: A Prospective Study in Project Viva. Epigenomics. 2019;11(12):1413–1427. doi:10.2217/epi-2019-0040

40. Alfano R, Guida F, Galobardes B, et al. Socioeconomic position during pregnancy and DNA methylation signatures at three stages across early life: epigenome-wide association studies in the ALSPAC birth cohort. Int J Epidemiol. 2019;48(1):30–44. doi:10.1093/ije/dyy259

41. Dunn EC, Soare TW, Zhu Y, et al. Sensitive Periods for the Effect of Childhood Adversity on DNA Methylation: Results From a Prospective, Longitudinal Study. Biol Psychiatry. 2019;85(10):838–849. doi:10.1016/j.biopsych.2018.12.023

42. Provençal N, Arloth J, Cattaneo A, et al. Glucocorticoid exposure during hippocampal neurogenesis primes future stress response by inducing changes in DNA methylation. Proc Natl Acad Sci. 2020;117(38):23280–23285. doi:10.1073/pnas.1820842116

43. Gluckman PD, Hanson MA, Buklijas T. A conceptual framework for the developmental origins of health and disease. J Dev Orig Health Dis. 2010;1(1):6–18. doi:10.1017/S2040174409990171

44. Hetz C, Martinon F, Rodriguez D, Glimcher LH. The Unfolded Protein Response: Integrating Stress Signals Through the Stress Sensor IRE1α. Physiol Rev. 2011;91(4):1219–1243. doi:10.1152/physrev.00001.2011

45. Riaz TA, Junjappa RP, Handigund M, Ferdous J, Kim HR, Chae HJ. Role of Endoplasmic Reticulum Stress Sensor IRE1α in Cellular Physiology, Calcium, ROS Signaling, and Metaflammation. Cells. 2020;9(5):1160. doi:10.3390/cells9051160

46. Magg V, Manetto A, Kopp K, et al. Turnover of PPP1R15A mRNA encoding GADD34 controls responsiveness and adaptation to cellular stress. Cell Rep. 2024;43(4). doi:10.1016/j.celrep.2024.114069

47. Oliveira MM, Mohamed M, Elder MK, et al. The integrated stress response effector GADD34 is repurposed by neurons to promote stimulus-induced translation. Cell Rep. 2024;43(2). doi:10.1016/j.celrep.2023.113670

48. Hetz C, Saxena S. ER stress and the unfolded protein response in neurodegeneration. Nat Rev Neurol. 2017;13(8):477–491. doi:10.1038/nrneurol.2017.99

49. Kim P. Understanding the Unfolded Protein Response (UPR) Pathway: Insights into Neuropsychiatric Disorders and Therapeutic Potentials. Biomol Ther. 2024;32(2):183–191. doi:10.4062/biomolther.2023.181

50. Solarz-Andrzejewska A, Majcher-Maślanka I, Kryst J, Chocyk A. Modulation of the endoplasmic reticulum stress and unfolded protein response mitigates the behavioral effects of early-life stress. Pharmacol Rep. 2023;75(2):293–319. doi:10.1007/s43440-023-00456-6

51. Liu Y, Pan Z, Shi J, Jiang M, Li J. A novel heterozygous mutation of BCL11B gene causes neurodevelopmental disorder and middle type hypospadias in a Chinese boy with 5 years follow-up. Neurogenetics. 2026;27(1):13. doi:10.1007/s10048-026-00880-9

52. Seigfried FA, Britsch S. The Role of Bcl11 Transcription Factors in Neurodevelopmental Disorders. Biology. 2024;13(2):126. doi:10.3390/biology13020126

53. Lussier AA, Smith BJ, Fisher J, et al. DNA methylation mediates the link between adversity and depressive symptoms. Nat Ment Health. 2024;2(12):1476–1485. doi:10.1038/s44220-024-00345-8

54. Li S, Spitz N, Ghantous A, et al. A Pregnancy and Childhood Epigenetics Consortium (PACE) meta-analysis highlights potential relationships between birth order and neonatal blood DNA methylation. Commun Biol. 2024;7(1):66. doi:10.1038/s42003-023-05698-x

55. Volkow ND, Gordon JA, Bianchi DW, et al. The HEALthy Brain and Child Development Study (HBCD): NIH collaboration to understand the impacts of prenatal and early life experiences on brain development. Dev Cogn Neurosci. 2024;69:101423. doi:10.1016/j.dcn.2024.101423

56. Schuurmans IK, Mulder RH, Baltramonaityte V, et al. Consortium profile: the methylation, imaging and NeuroDevelopment (MIND) consortium. Mol Psychiatry. 2026;31(2):1177–1189. doi:10.1038/s41380-025-03203-w

57. Álvarez-Mejía D, Rodas JA, Leon-Rojas JE. From Womb to Mind: Prenatal Epigenetic Influences on Mental Health Disorders. Int J Mol Sci. 2025;26(13):6096. doi:10.3390/ijms26136096

58. Fraser A, Macdonald-Wallis C, Tilling K, et al. Cohort Profile: The Avon Longitudinal Study of Parents and Children: ALSPAC mothers cohort. Int J Epidemiol. 2013;42(1):97–110. doi:10.1093/ije/dys066

59. Tobi EW, Heuvel J van den, Zwaan BJ, Lumey LH, Heijmans BT, Uller T. Selective Survival of Embryos Can Explain DNA Methylation Signatures of Adverse Prenatal Environments. Cell Rep. 2018;25(10):2660–2667.e4. doi:10.1016/j.celrep.2018.11.023

60. van den Oord CLJD, Copeland WE, Zhao M, Xie LY, Aberg KA, van den Oord EJCG. DNA methylation signatures of childhood trauma predict psychiatric disorders and other adverse outcomes 17 years after exposure. Mol Psychiatry. 2022;27(8):3367–3373. doi:10.1038/s41380-022-01597-5

61. Radtke KM, Schauer M, Gunter HM, et al. Epigenetic modifications of the glucocorticoid receptor gene are associated with the vulnerability to psychopathology in childhood maltreatment. Transl Psychiatry. 2015;5(5):e571–e571. doi:10.1038/tp.2015.63

